# Atypical *Salmonella enterica* serovars in murine and human infection models: Is it time to reassess our approach to the study of salmonellosis?

**DOI:** 10.1101/058610

**Authors:** Daniel Hurley, Maria Hoffmann, Tim Muruvanda, Marc W. Allard, Eric W. Brown, Marta Martins, Séamus Fanning

**Affiliations:** UCD-Centre for Food Safety, School of Public Health, Physiotherapy and Sports Science, University College Dublin, Belfield, Dublin D04 N2E5, Ireland; Center for Food Safety and Nutrition, Division of Microbiology, Office of Regulatory Science, U.S. Food and Drug Administration, College Park, Maryland 20740, USA

## Abstract

Nontyphoidal *Salmonella* species are globally disseminated pathogens and the predominant cause of gastroenteritis. The pathogenesis of salmonellosis has been extensively studied using *in vivo* murine models and cell lines typically challenged with *Salmonella* Typhimurium. Although serovars Enteritidis and Typhimurium are responsible for the most of human infections reported to the CDC, several other serovars also contribute to clinical cases of salmonellosis. Despite their epidemiological importance, little is known about their infection phenotypes. Here, we report the virulence characteristics and genomes of 10 atypical *S. enterica* serovars linked to multistate foodborne outbreaks in the United States. We show that the murine RAW 264.7 macrophage model of infection is unsuitable for inferring human relevant differences in nontyphoidal *Salmonella* infections whereas differentiated human THP-1 macrophages allowed these isolates to be further characterised in a more relevant, human context.

---

*Salmonella* is a zoonotic pathogen responsible for illnesses on a global scale and poses a significant burden to public health^1^. Invasive salmonellae, such as the host-restricted *Salmonella enterica* (*S. enterica*) serovars Typhi and Paratyphi cause fever in humans killing nearly 217,000 people worldwide^2^. Infection with nontyphoidal *Salmonella* (NTS) such as *S. enterica* serovars Enteritidis and Typhimurium results in bacteraemia and gastroenteritis and is estimated to cause 155,000 deaths annually^3^.

*Salmonella* are non-fastidious bacteria that can survive outside the host in a range of food matrices, low-moisture conditions as well as food processing environments. In humans, infection commonly occurs following the ingestion of contaminated food or water. Sources for contamination vary and can range from the presence of the bacterium on raw produce to the shedding of *Salmonella* in the faecal and urinary excretions of reservoir animals^4,5^.

Upon entering the host, salmonellae are challenged by a series of adverse conditions including the low pH environment of the stomach, the membrane disrupting properties of bile in the small intestine and a battery of phagocytic host immune cells such as macrophages^67^. Deployment of a <u>T</u>ype <u>III</u> Secretion <u>S</u>ystem (T3SS) apparatus is fundamental to the pathogenesis of *Salmonella* and enables the bacterium to translocate effector proteins into the host cell cytoplasm^8^. The acquisition of <u>S</u>almonella <u>P</u>athogenicity <u>i</u>sland (SPI) encoded virulence factors *via* horizontal gene transfer followed by evolution has enabled this microorganism to exploit a privileged replicative niche, avoiding the host innate immune system within intracellular vesicles called *<u>S</u>almonella*-<u>C</u>ontaining <u>V</u>acuole (SCV)^9,10^. The protection afforded by the SCV allows *Salmonella* to thrive and sets in motion a cycle of infection whereby the bacterium can proliferate and basolaterally reinfect epithelial cells as well as become engulfed by additional, localised phagocytic cells.

The importance of *Salmonella* in both clinical and public health settings has fuelled research into the virulence mechanisms and associated pathogenicity of this bacterium^11,12^. Many of these studies use *S. Typhimurium* infection in mice as a model for typhoid fever due to the similar pathology observed including intestinal and extraintestinal lesions that frequently occur in human hosts infected with typhoidal serovars^13^. The *in vitro* transcriptome of S. Typhimurium has been characterised extensively by sequencing-based methods^14,15^. Similarly, the *ex vivo* response of *S.* Typhi and *S.* Typhimurium during infection of macrophages has also been elucidated^16–18^.

Although *S*. Typhimurium infection of mice results in symptoms mimicking human typhoid fever, this serovar, along with S. Enteritidis is predominantly associated with gastroenteritis in humans. Despite the fact that over 2,610 serovars of *Salmonella* have been reported to date, few studies are available to describe the infection arising from other *S. enterica* serovars^19–22^.

In this study, 10 atypical *S. enterica* serovars cultured from multistate foodborne outbreaks in the U.S. were characterised both pheno- and genotypically. Differences in the nature of the infection process and the underlying virulence determinants were noted. We identified factors that further explain the ability of these atypical *S. enterica* serovars to cause foodborne outbreaks.

## Results

### *In vitro* characterisation of the isolates

To simulate the host-specified challenges ingested salmonellae encounter along the alimentary canal during infection in an *in vitro* laboratory environment, isolates (Table 1) were characterised for their acid tolerance and swim/swarm motility. In addition, differences in susceptibility to salts of the main acid constituents of bile were determined by <u>M</u>inimum <u>I</u>nhibitory <u>C</u>oncentration (MIC) and <u>M</u>inimum <u>B</u>actericidal <u>C</u>oncentration (MBC) assays.

**Table 1.** https://dx.doi.org/10.6084/m9.figshare.3364726.V1

| **S. enterica serovar** | **Strain** | **Outbreak** | **Location** | **Year** | **Isolation Source** |
| --- | --- | --- | --- | --- | --- |
| Anatum | CFSAN003959 | Papaya | Mexico | 2012 | Papaya |
| Bareilly | CFSAN001111 | Tuna | India | 2012 | Tuna scrape, frozen |
| Cubana | CFSAN002050 | Alfalfa sprouts | USA - Arizona | 2012 | Alfalfa sprouts |
| Heidelberg | CFSAN002063 | Chicken | USA - Washington | 2012 | Clinical |
| Javiana | CFSAN000905 | Green onion | Mexico | 2009 | Canal water |
| Montevideo | CFSAN000264 | Black pepper | USA - Rhode Island | 2010 | Black pepper |
| Newport | CFSAN003345 | Eastern shore sampling | USA - Virginia | 2011 | Goose faeces |
| Saintpaul | CFSAN004090 | Jalapeño/serrano pepper | Mexico | 2008 | Jalapeño pepper |
| Tennessee | CFSAN001387 | Peanut butter | USA - Georgia | 2007 | Peanut butter |
| **Typhimurium** | **14028S** |  |  |  |  |
| **Typhimurium** | **ST4/74** |  |  |  |  |
| Weltevreden | CFSAN001415 | Prison tuna | USA - Virginia | 2005 | Tuna |

The acid tolerance response of *Salmonella* is a complex defence mechanism employed by the pathogen to defend against the acid shock experienced in the stomach^6^. Increased numbers for the majority of isolates were recorded following 1 and 2 hours of growth at pH 2.5 before significant reductions in viable numbers occurred at 4 hours (Supplementary Fig. S1). Motility has been suggested as a virulence determinant with respect to invasion with notable non-motile and/or host adapted exceptions^23–26^. After 24 hours of incubation at 21 °C, *S*. Bareilly CFSAN001111 and *S*. Javiana CFSAN000905 exhibited a reduced swim phenotype when compared to *S*. Typhimurium ST4/74 (P < 0.001) whereas S. Newport CFSAN003345 and *S*. Saintpaul CFSAN004090 exhibited an increased swim phenotype (P < 0.001). At 37 °C, *S*. Anatum CFSAN003959 and *S*. Tennessee CFSAN001387 exhibited a reduced swim phenotype after 8 and 24 hours when compared to S. Typhimurium ST4/74 (P < 0.001) (Supplementary Fig. S2). In the absence of glucose as a carbon source, no swarm motility was observed for any of the isolates regardless of temperature or incubation time (Supplementary 5 Fig. S2). During digestion, contraction of the gall bladder releases bile into the small intestine. *Salmonella* Weltevreden CFSAN001415 showed a two-fold difference in susceptibility to sodium deoxycholate (DOC) in comparison to S. Typhimurium ST4/74 although the MBC was comparable to the other isolates (Supplementary Table S1).

### Intracellular survival of atypical *S. enterica* serovars in murine and human macrophages

*Salmonella* Typhimurium pathogenesis has been extensively studied *in vivo* using murine models and *ex vivo* using murine cell lines (such as J774.2 and RAW 264.7). However, the differences in the ability of *Salmonella* to survive and replicate within human macrophages is currently not well described^27^. Differentiated human monocyte cell lines (including THP-1 and U937) have been used to explore the replication of *S. enterica* serovars Enteritidis and Typhimurium. Few studies describing the bacterial replication in human macrophages of atypical NTS serovars have been reported despite their epidemiological importance and their contributions to clinical cases of salmonellosis^27,28^.

To study the ability of the isolates to survive phagocytosis, infections were performed using murine RAW 264.7 and differentiated human THP-1 macrophages by the gentamicin protection assay adapted from protocols previously described^29–32^. *Salmonella* Typhimurium 14028S and ST4/74 were included as reference strains in all infection assays. Infections were carried out at a <u>M</u>ultiplicity <u>O</u>f <u>I</u>nfection (MOI) of 10:1. Viable internalised bacteria were enumerated at 2, 4, 8 and 24 <u>H</u>ours <u>P</u>ost <u>I</u>nfection (HPI) in RAW 264.7 and 2, 4, 8, 24 and 168 HPI in THP-1 macrophages.

Of the 10 atypical serovars tested in this study, all isolates were found to persist within RAW 264.7 macrophages for 24 HPI with many of these increasing in number over the course of the infection. In the case of *S*. Weltevreden CFSAN001415, a 1-Log_10_ decrease in intracellular bacteria between 2 and 24 HPI was recorded though this isolate was still recoverable. In contrast, *S*. Tennessee CFSAN001387 exhibited a 1-Log_10_ increase in intracellular bacteria between 2 and 24 HPI. Similar numbers of viable bacteria were recorded for *S*. Typhimurium 14028S and ST4/74 (Fig. 1a).

**Fig. 1:**
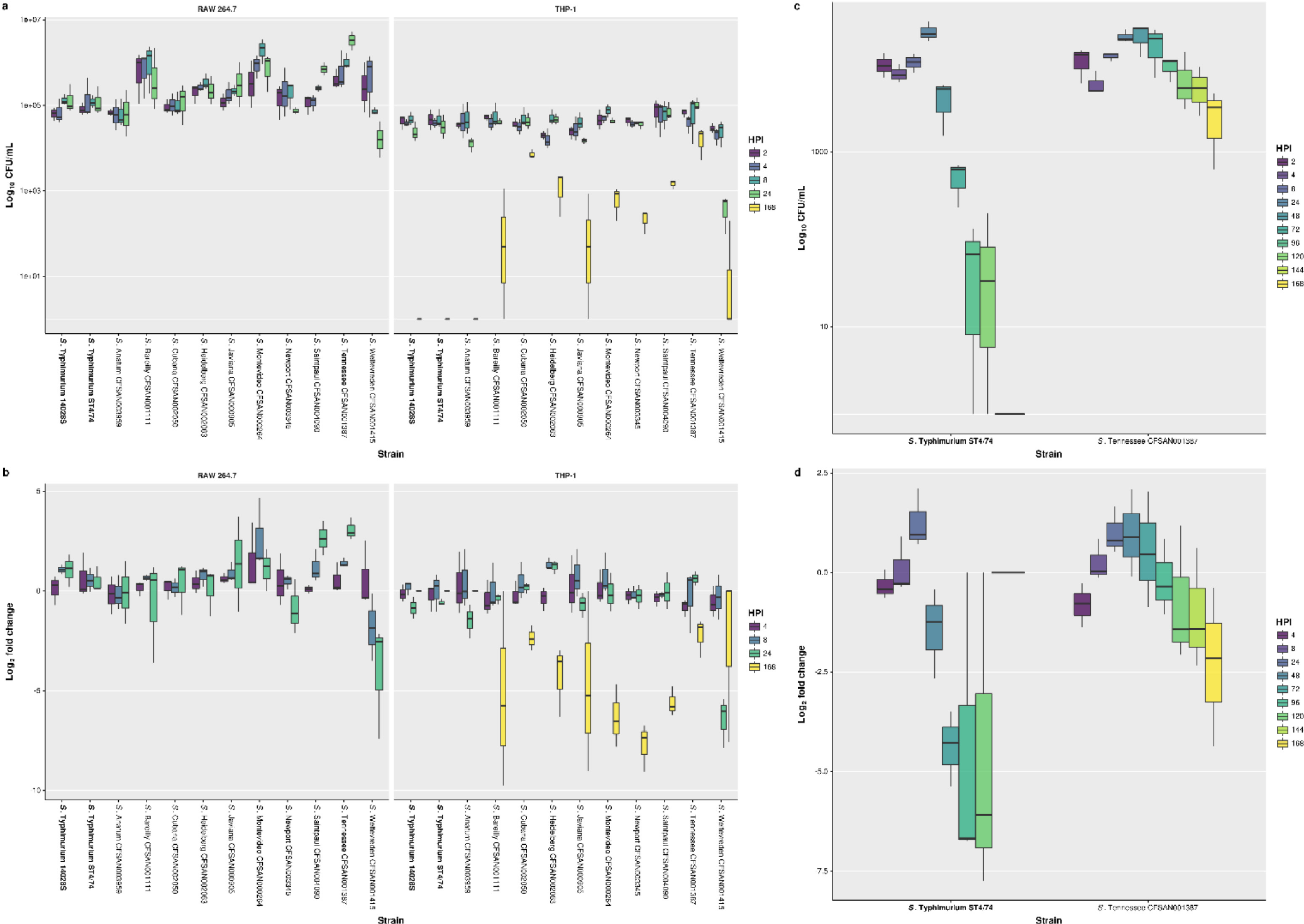
Survival of isolates following phagocytosis by RAW 264.7 and THP-1 macrophages assessed by gentamicin protection assay. **A)** Survival of isolates following phagocytosis by RAW 264.7 macrophages at 2, 4, 8 and 24 HPI and THP-1 macrophages at 2, 4, 8, 24 and 168 HPI reported as Log10 CFU/mL. **B)** Survival of isolates following phagocytosis by RAW 264.7 macrophages at 4, 8 and 24 HPI and THP-1 macrophages at 4, 8, 24 and 168 HPI reported as Log2 fold change comparing to 2 HPI. Results for isolates in the study correspond to the mean of three independent assays (*n* = 3) with duplicate technical replicates. Results for reference strains S. Typhimurium 14028Sand ST4/74 correspond to the mean of six independent assays (*n* = 6) with duplicate technical replicates. **C)** Survival of S. Tennessee CFSAN001387 in comparison with S. Typhimurium ST4/74 following phagocytosis by THP-1 macrophages at 2, 4, 8, 24, 48, 72, 96, 120, 144 and 168 HPI reported as Log10 CFU/mL. **D)** Survival of S. Tennessee CFSAN001387 in comparison with *S*. Typhimurium ST4/74 following phagocytosis by THP-1 macrophages at 4, 8, 24, 48, 72, 96, 120, 144 and 168 HPI reported as Log2 fold change comparing to 2 HPI. Results correspond to the mean of three independent assays (*n* = 3) with triplicate technical replicates. The lower and upper hinges correspond to the 25th and 75th percentiles with the whiskers extending ± 1.5 times the range between first and third quartiles. https://dx.doi.org/10.6084/m9.figshare.3364726.V1

The highest and lowest mean <u>C</u>olony <u>F</u>orming <u>U</u>nits (CFU)/mL for all 10 atypical serovars are shown (Supplementary Table S2). When the Log2 fold change was calculated, *S*. Weltevreden CFSAN001415 recorded the largest decrease at 8 and 24 HPI (Fig. 1b). Of note, *S*. Bareilly CFSAN001111 showed an unusual infection profile in that it had the highest mean CFU/mL at 2 HPI, possibly indicative of increased invasiveness, despite normalisation of the infections by centrifuging the cell culture plates. The latter observation was maintained for all subsequent time points. However, the mean CFU/mL at 4, 8 and 24 HPI for *S*. Bareilly CFSAN001111 were lower than other isolates.

When compared to *S*. Typhimurium ST4/74, significant differences were observed in the infection profile of specific isolates at individual time points (Supplementary Table S3). Time courses longer than 24 hours were not possible in RAW 264.7 as this resulted in cell death or proliferation of the macrophages themselves which would skew the MOI. Although differences between these isolates were observed in RAW 264.7 macrophages, the inability to consistently extend the time course of the assay beyond 24 HPI limited the utility of this *ex vivo* murinemodel to discern any further differences in the ability of the isolates to survive and persist following phagocytosis. To mitigate this limitation and differentiate the isolates in a human relevant context, additional assays were carried out in differentiated THP-1 macrophages.

Of the 10 atypical serovars tested in this study, all persisted within THP-1 macrophages for 24 HPI with the majority exhibiting no significant changes in viable numbers up to this time point. As with RAW 264.7 macrophages, the only exception noted when infecting THP-1 macrophages was recorded for S. Weltevreden CFSAN001415 which exhibited a 1-Log_10_ decrease in mean CFU/mL between 2 and 24 HPI but remained recoverable. Upon extending this assay beyond 24 hours to 168 HPI, equivalent to 7 days, all serovars with the exception of S. Anatum CFSAN003959 and both S. Typhimurium 14028S and ST4/74 reference strains were recoverable with the majority exhibiting a 2-Log_10_ decrease in intracellular bacteria. Exceptions to this were noted for S. Cubana CFSAN002050, S. Heidelberg CFSAN002063 and S. Tennessee CFSAN001387 which showed a 1-Log_10_ decrease in intracellular bacteria between 2 and 168 HPI with S. Tennessee CFSAN001387 displaying the smallest decrease of all isolates. Conversely, S. Weltevreden CFSAN001415 demonstrated the largest decrease in bacterial cell numbers at 168 HPI but unlike S. Anatum CFSAN003959 and both S. Typhimurium 14028S and ST4/74 reference strains, it could still be recovered at the end of the assay (Fig. 1a).

The highest and lowest mean CFU/mL among the 10 study isolates are shown (Supplementary Table S2). In THP-1 macrophages, S. Weltevreden CFSAN001415 had the lowest mean CFU/mL at 24 and 168 HPI of the recoverable isolates as observed in the Log_2_ fold change (Fig. 1b). As observed in RAW 264.7 macrophages, S. Tennessee CFSAN001387 had the highest mean CFU/mL at 24 HPI as well as 168 HPI in THP-1 macrophages. When compared to S. Typhimurium ST4/74, significant differences were observed in the infection profiles of specific isolates at individual time points (Supplementary Table S2).

Fewer significant differences were observed between isolates when compared to S. Typhimurium ST4/74 infection in human THP-1 macrophages versus murine RAW 264.7 macrophages highlighting the potential unsuitability of the murine model for inferring human relevant differences between isolates in NTS infection. Overall, the viable intracellular bacteria recorded at later time points, including 24 HPI, was significantly higher in RAW 264.7 than THP-1. Bacterial cell numbers reached as high as 1 × 10^7^ mean CFU/mL in RAW 264.7 for some atypical serovars compared with values that did not exceed 1 × 10^5^ mean CFU/mL in THP-1 for all isolates. This 2-Log_10_ difference supports our observations showing the inability of RAW 264.7 to clear infecting bacteria compared with THP-1 macrophages.

The viability of both murine and human macrophages following infection with each of the selected bacterial isolates was measured using colorimetric assays to measure extracellular <u>G</u>lucose <u>6</u>-<u>P</u>hosphate (G6P) and <u>L</u>actate <u>D</u>eHydrogenase (LDH) activities compared with uninfected control macrophages. No significant differences were observed in host cell viability following a MOI of 10:1 (Supplementary Fig. S4 and Supplementary Fig. S5). This is in agreement with recent studies that used flow cytometry based techniques to quantify apoptosis in macrophages infected with different *Salmonella* strains^27,33^.

As S. Tennessee CFSAN001387 was noted to be the most proliferative isolate in THP-1 macrophages, exhibiting the lowest reduction in viable intracellular bacteria between 2 and 168 HPI, further assays were performed to directly compare to S. Typhimurium ST4/74 with CFU being enumerated at 2, 4, 8, 24, 48, 72, 96, 120, 144 and 168 HPI. This was done to determine when S. Typhimurium ST4/74 was no longer recoverable compared with S. Tennessee CFSAN001387. *Salmonella* Typhimurium ST4/74 exhibited an overall 1-Log10 reduction between 2 and 48 HPI with an additional 2-Log_10_ reduction in viable bacteria by 72 hours. After 96 and 120 HPI, S. Typhimurium ST4/74 was barely detected and was unrecoverable at 144 and 168 HPI. In comparison, S. Tennessee CFSAN001387 exhibited an overall 1-Log10 reduction between 2 and 168 HPI, similar to the previous infections (Fig. 1c). Differences between these two isolates were significant at multiple individual time points (Supplementary Table S4).

### Host response to atypical S. *enterica* infection by murine and human macrophages

As there were notable differences exhibited by these *Salmonella* serovars in their ability to replicate and survive within murine and human macrophages, the host response to infection was investigated by quantifying proinflammatory cytokine and infection relevant chemokine release using standard immunoassay protocols.

In RAW 264.7, increased CCL2 and CCL3 chemokine release was observed at 4, 8 and 24 HPI with S. Tennessee CFSAN001387 and S. Weltevreden CFSAN001415 when compared to uninfected control macrophages. When comparing infection with these 10 atypical serovars (Table 1) to S. Typhimurium ST4/74, increased proinflammatory cytokine (including IL6, IL10 and TNF) and chemokine (including CCL2, CCL3 and CXCL10) release was observed (Fig. 2) (Supplementary Table S5).

**Fig. 2:**
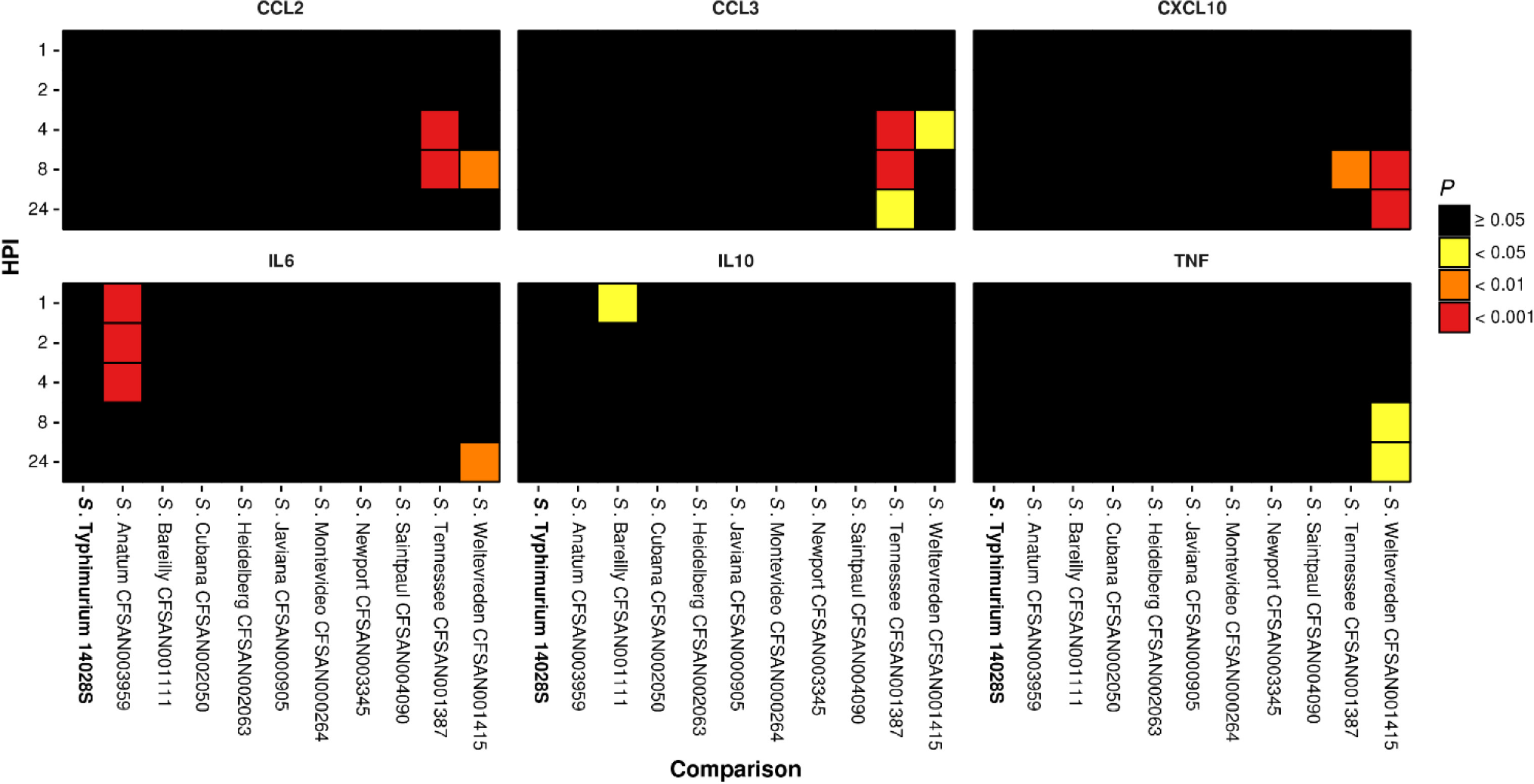
Significant RAW 264.7 chemokine and proinflammatory cytokine release following infection with isolates when compared to S. Typhimurium ST4/74 infection. Increased release of chemokine and proinflammatory cytokine proteins by RAW 264.7 macrophages at 1, 2, 4, 8 and 24 HPI with isolates when compared to S. Typhimurium ST4/74 as determined by one-way ANOVA. Values correspond to adjusted probability (P) as determined by *post hoc* analysis of significance using Tukey’s range test. Differences were deemed significant by arbitrary cut-offs at < 0.05, 0.01 and 0.001. https://dx.doi.org/10.6084/m9.figshare.3364726.V1

In THP-1, increased proinflammatory cytokine (including CXCL8, IL1B, IL6 and TNF), cytokine (including CSF2, IL1A, IL12B and VEGFA) and chemokine (including CCL2, CCL3, CCL4 and CXCL10) release was recorded at 8, 24 and 168 HPI across a selection of these isolates in comparison with uninfected control macrophages. As with RAW 264.7, when comparing infection with the 10 atypical serovars in this study to S. Typhimurium ST4/74 in THP-1, significant increased proinflammatory cytokine and chemokine release was observed, further differentiating the isolates by the innate host response (Fig. 3) (Supplementary Table S5).

**Fig. 3:**
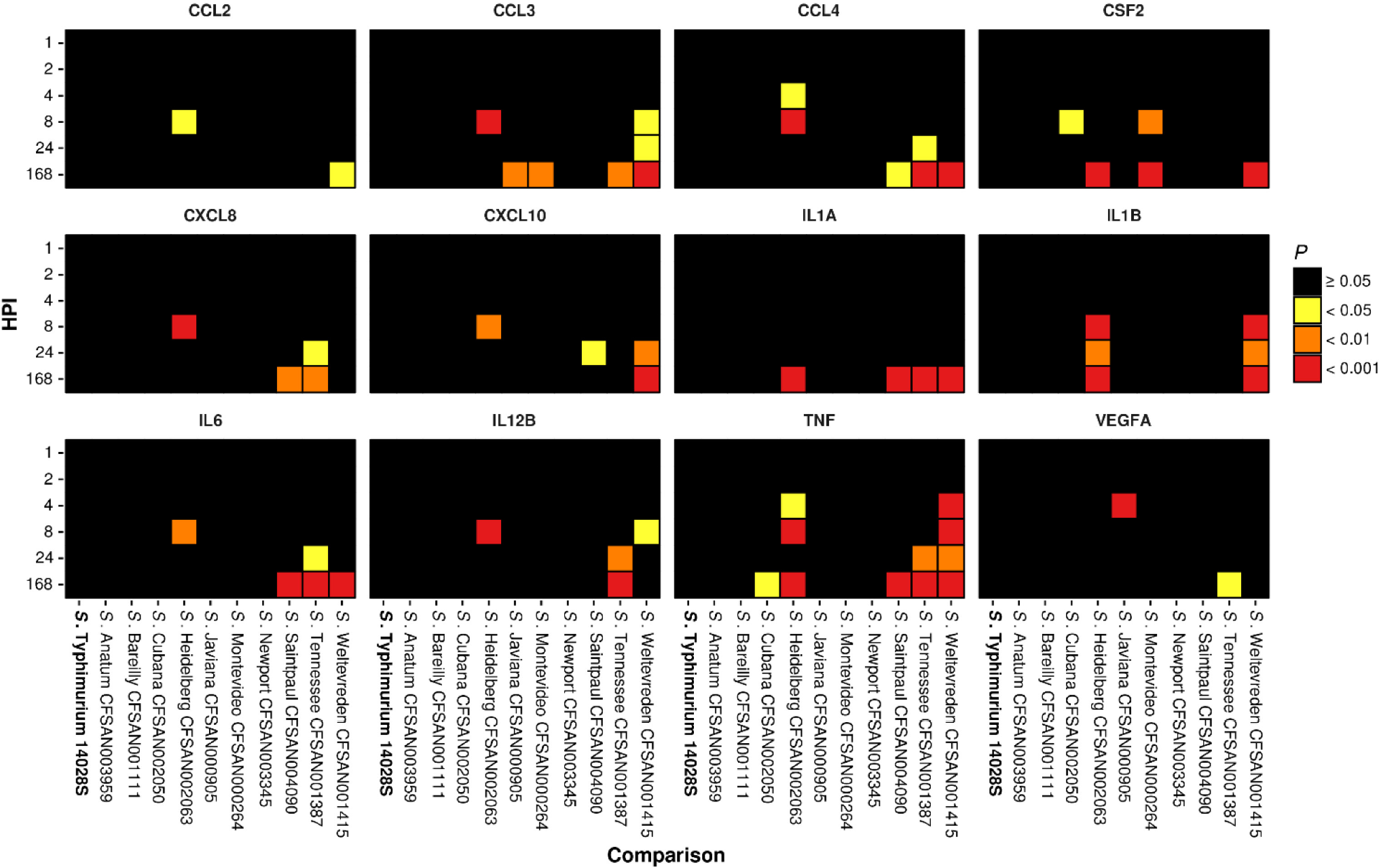
Significant THP-1 chemokine, cytokine and proinflammatory cytokine release following infection with isolates when compared to S. Typhimurium ST4/74 infection. Increased release of chemokine, cytokine and proinflammatory cytokine proteins by THP-1 macrophages at 1, 2, 4, 8, 24 and 168 HPI with isolates when compared to S. Typhimurium ST4/74 as determined by one-way ANOVA. Values correspond to adjusted probability (P) as determined by *post hoc* analysis of significance using Tukey’s range test. Differences were deemed significant by arbitrary cut-offs at < 0.05, 0.01 and 0.001. https://dx.doi.org/10.6084/m9.figshare.3364726.V1

*Salmonella* Heidelberg CFSAN002063 stimulated the release of CCL3, CSF2, CXCL8, CXCL10, IL1A, IL1B, IL6, IL12 and TNF to levels in excess of those observed for S. Typhimurium ST4/74, particularly at 8 HPI and from 4 to 168 HPI with respect to TNF (Supplementary Fig. S7). In the gentamicin protection assays it was noted that although both S. Tennessee CFSAN001387 and S. Weltevreden CFSAN001415 could be recovered at 168 HPI from THP-1 macrophages, these two isolates exhibited different infection profiles. The former displayed the smallest reduction in viable intracellular bacteria over the time course of the infection whereas the latter displayed the largest reduction in viable numbers. This can be accounted for in the overlapping yet contrasting cytokine profile of THP-1 macrophages following infection with these isolates. *Salmonella* Tennessee CFSAN001387 and S. Weltevreden CFSAN001415 stimulated the release of CCL3, CCL4, IL1A, IL6, IL12B and TNF to levels in excess of those observed for S. Typhimurium ST4/74 (Fig. 4a, Fig. 4b and Fig. 4c). In addition, S. Weltevreden CFSAN001415 stimulated the release of CCL2, CXCL10, CSF2 and IL1B triggering a broader response in comparison to that observed for S. Tennessee CFSAN001387 (Fig. 4a, Fig. 4b and Fig. 4c).

**Fig. 4:**
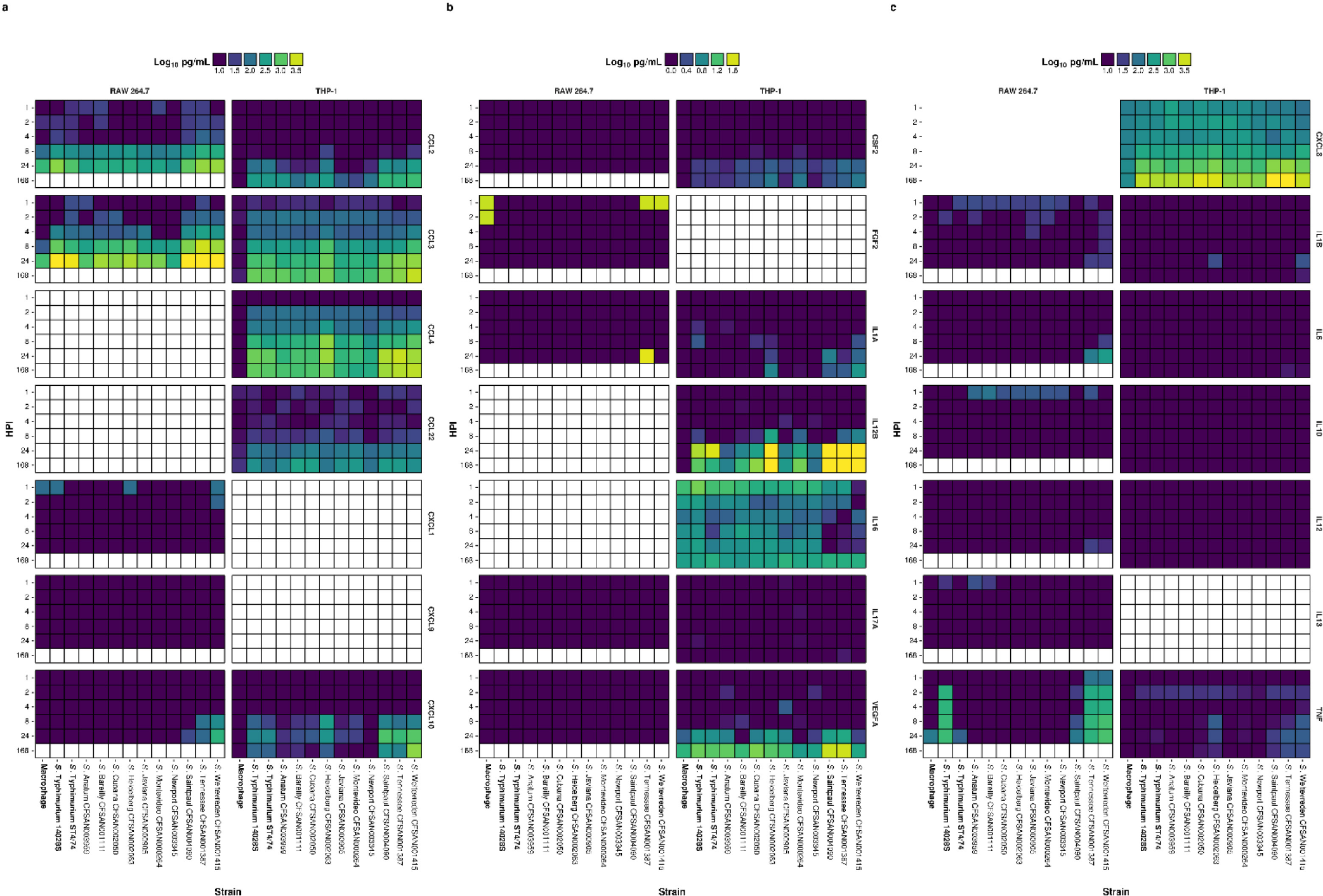
Chemokine, cytokine and proinflammatory cytokine release from infected RAW 264.7 and THP-1 macrophages. Release of chemokine (A), cytokine (B) and proinflammatory cytokine (C) targets by RAW 264.7 and THP-1 macrophages following infection with isolates reported as Log_10_ pg/mL. Cells coloured white correspond to time points not assayed with respect to protein or cell line. https://dx.doi.org/10.6084/m9.figshare.3364726.V1

Overall, the human macrophages mounted a much greater proinflammatory response to infection in comparison to the murine model (Fig. 4c). A homolog of the human *CXCL8* gene is absent in mice. However, the murine *Cxcll* gene codes for a functionally homologous protein^34^. The latter was not released from infected RAW 264.7 to the levels observed for CXCL8 in THP-1 in this study. As all infections were carried out in pure macrophage cultures, the true effect of the observed chemokine release cannot be fully appreciated in this experimental model as the activation and/or recruitment of other phagocytic cells in a coculture population or an *in vivo* environment would have an impact on bacterial survival.

### Distribution and similarity of SPI proteins from atypical S. *enterica* serovars in comparison to S. Typhimurium ST4/74

Whole genome sequencing was used to facilitate a comparative analysis of all 10 atypical serovars to elucidate the genetic variation among them with particular focus on virulence determinants and SPI loci gene content^35,36^. Sequencing was performed using the Illumina MiSeq and Pacific Biosciences RS II sequencing platforms. *De novo* assemblies were performed on these data and the N50 length for Illumina sequenced assemblies ranged from 403 to 763 kbp with an average N50 length of 586 kbp (Supplementary Table S6). These genome sequences were used to determine the relationships between the isolates to identify key differences that may explain the phenotypes observed in the previous experiments. Genetic diversity was characterised by amino acid sequence similarity that identified highly conserved SPI regions between these isolates despite the broad range of serovars in addition to distinct variable regions and/or absence of key effector proteins in specific isolates (Fig. 5 and Supplementary Fig. S8).

**Fig. 5:**
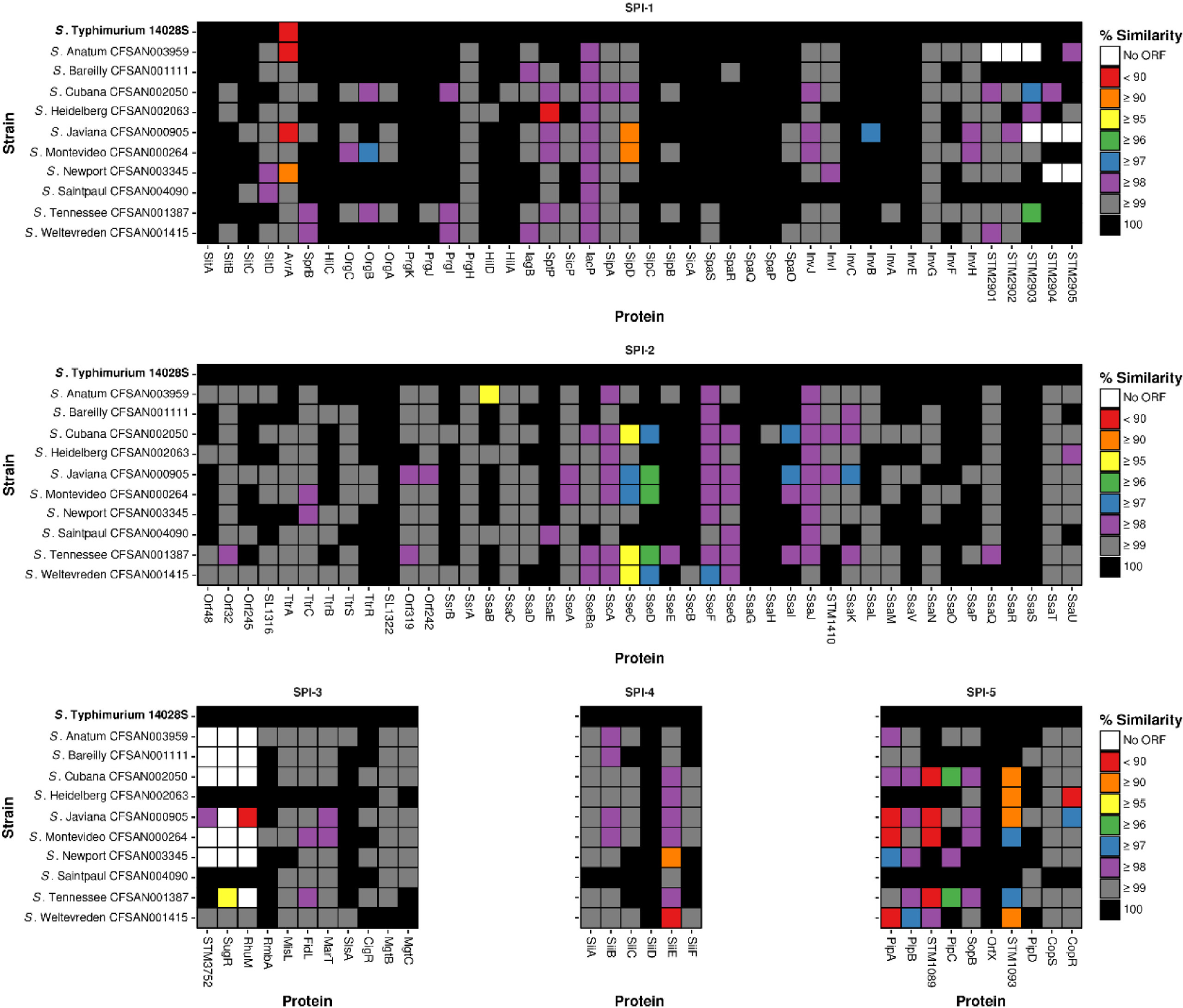
Sequence variability within SPI-1 to SPI-5. The amino acid sequence similarity for SPI gene content of each isolate included in this study was compared with S. Typhimurium ST4/74. https://dx.doi.org/10.6084/m9.figshare.3364726.V1

SPI-1 and its associated T3SS have been extensively implicated in *Salmonella* virulence and the ability of this pathogen to invade host eukaryotic cells, trigger inflammation and transport effector proteins^37,38^. Sequence variation at the amino acid level in comparison to S. Typhimurium ST4/74 was greatest in AvrA, OrgB, SptP, SipD, InvB and SL2883 proteins. Several SPI-1 encoded genes were absent in many of the atypical serovars but present in S. Typhimurium ST4/74 though not yet fully characterised (Fig. 5).

The AvrA protein has been previously shown to be crucial, playing a role in the inhibition of the antiapoptotic NF-KB pathway^39^. In S. Typhimurium 14028S, S. Anatum CFSAN003959, S. Javiana CFSAN000905 and S. Newport CFSAN003345, AvrA showed differences at the amino acid sequence level which may affect its ability to function as a protease^40^. SptP is a tyrosine phosphatase involved in the inhibition Raf activation and the subsequent MAP kinase pathway^41^. Its identification here is consistent with the phenotype observed for S. Heidelberg CFSAN002063 that stimulated high levels of TNF release at 4, 8, 24 and 168 HPI in THP-1 macrophages. A number of proteins, including STM2901, STM2902, STM2903, SL2883, STM2904 and STM2905 that are SPI-1 associated in S. Typhimurium ST4/74 showed mixed
conservation, either being highly similar in the majority of isolates or were unidentifiable in S. Anatum CFSAN003959, S. Javiana CFSAN000905 and S. Newport CFSAN003345.

SPI-2 and its associated T3SS contributes to the ability of *Salmonella* to translocate effectors across the membrane of the SCV when the bacterium is internalised in epithelial cells and macrophages^42,43^. The integral function of SPI-2 for intracellular survival can be observed by the degree of amino acid sequence conservation across all atypical serovars (Fig. 5). As these isolates were implicated in multistate foodborne outbreaks and, as demonstrated above, capable of surviving within both RAW 264.7 and THP-1 macrophages, this observation is consistent with the expressed phenotype and raises the question as to whether or not potential differences in the expression of some/all of these genes may further explain the differences shown in intracellular survival. Sequence variation at the amino acid level in comparison to S. Typhimurium ST4/74 was greatest in SsaB, SseB, SseC and SseD. Loss of the effector protein SsaB (SpiC) has been shown to promote defective virulence phenotypes due to an inability to translocate all SPI-2 effectors^44^. This may explain the observed infection phenotype in S. Anatum CFSAN003959, the only isolate among the 10 atypical serovars studied to be unrecoverable at 168 HPI in THP-1 macrophages, as it had the lowest amino acid sequence similarity for SsaB.

SseBCD proteins function as a translocon that facilitate the secretion of effector proteins by intracellular *Salmonella*^45^. Both S. Cubana CFSAN002050 and S. Tennessee CFSAN001387 display differences in all three SseBCD proteins with S. Tennessee CFSAN001387 being least similar at the amino acid sequence level when compared to S. Typhimurium ST4/74.

*Salmonella* Tennessee CFSAN001387 was shown in this study to survive within THP-1 macrophages significantly better in comparison to S. Typhimurium ST4/74.

High levels of similarity to S. Typhimurium ST4/74 were observed for SPI-3 across the isolates with the exception of the complete loss or major differences in STM3752, SugR and RhuM as reported previously^35^. SPI-4 and its associated <u>T</u>ype <u>I</u> <u>S</u>ecretion <u>S</u>ystem (T1SS) have been implicated in adhesion, contributing to intestinal inflammation in animal models^46,47^. *Salmonella* Newport CFSAN003345 and S. Weltevreden CFSAN001415 exhibited low levels of amino acid sequence similarity to S. Typhimurium ST4/74 with respect to SiiE which has been shown to be important for persistent infection in macrophages^48^. SPI-5 encodes many SPI-1 and SPI-2 T3SS targeted effector proteins. Sequence variation was observed for CopR, PipA, STM1089, STM1093 in multiple isolates with mutations in PipA having previously been implicated in enteric salmonellosis^49^.

## Discussion

Studies aimed at elucidating the host response to NTS serovars including Enteritidis and Typhimurium have been facilitated by the availability of murine infection models. This study focussed on atypical serovars of this genus availing of isolates cultured from foodborne outbreaks, the majority of which are frequently listed in the top 20 serovars responsible for laboratory-confirmed human cases of salmonellosis as reported annually by the CDC^50^. These data identify key differences between isolates related to their ability to survive within macrophages and highlight the potential unsuitability of the widely used murine macrophage model to infer human relevant distinctions between isolates. We have shown that for NTS serovars, distinct differences in the inflammatory response of human macrophages can further differentiate these microorganisms in a manner that was not possible with murine macrophages (Fig. 3). Furthermore, the reference strains S. Typhimurium 14028S and ST4/74 that are often included in *in vitro* research emerged as the biological outliers in many respects with regards to their infection phenotype (Fig. 1). In the case of specific proinflammatory cytokines such as TNF, RAW 264.7 macrophages responded to infection with S. Typhimurium 14028S and ST4/74, S. Tennessee CFSAN001387 and S. Weltevreden CFSAN001415 whereas THP-1 macrophages displayed a broader proinflammatory response to the range of serovars studied (Fig. 4).

These data suggest more differences than previously acknowledged for S. *enterica* serovars with implications for public health. In agreement with this notion, key differences were identified in established virulence determinants of *Salmonella* such as SPI gene content bridging the gap 16between the observed phenotypes and the underlying genotypes. Further work will be required to understand the full scope of other potential targets within the genomes of these isolates such as the accessory gene content unique to individual strains, many of which are currently poorly characterised.

## Methods

### Bacterial isolates and culture methods

Environmental and food isolates were collected from different countries between 2005 and 2012 by the U.S. Food and Drug Administration inspections as part of compliance actions^51^. The clinical isolate was obtained from the Washington State Department of Health (Table 1). All *Salmonella* isolates were stored at −80 °C in <u>L</u>ysogeny <u>B</u>roth (LB) broth (Sigma-Aldrich) supplemented with 15 % [v/v] glycerol. Working cultures were prepared by streaking isolates and restreaking individual colonies onto <u>M</u>ueller-<u>H</u>inton (MH) agar (Sigma-Aldrich).

Individual colonies from restreaked isolates were used to inoculate 5 mL MH broth (Sigma-Aldrich) and grown overnight at 37 °C with orbital shaking at 200 RPM. Overnight cultures were then used in the subsequent experiments as detailed below.

### Acid resistance

The ability to survive in a low pH culture medium was assessed for all isolates as described previously^52^. Briefly, bacterial cultures were individually grown overnight (18 hours) without shaking in 5 mL buffered MH broth (MES hydrate, 2 % [w/v], pH 5) containing 0.4 % [w/v] glucose at 37 °C. A volume of 333 μL of overnight culture was centrifuged at 10,000 RPM for 10 minutes. The supernatant was removed and the cell pellet recovered and resuspended in 2 mL prewarmed, buffered MH broth (MES hydrate, 2 % [w/v], pH 2.5) containing 0.4 % [w/v] glucose and incubated at 37 °C in a 24 well plate. Time points were taken at 0 hours (before acidification), 1, 2 and 4 hours post-acidification. At each time point, 10 μL of the culture was diluted in 1,990 μL of maximum recovery diluent (MRD) medium (Oxoid) in a fresh 24 well plate and incubated for 30 minutes at room temperature to allow the bacteria to recover. Samples were then decimally diluted in phosphate buffered saline (PBS) (Sigma-Aldrich) and 100 μL aliquots of these dilutions were plated directly onto LB agar. Agar plates were incubated for 18 hours at 37 °C before enumeration of the CFU.

### Motility assays

Swim and swarm motility was assessed for all isolates as described previously^52^. Briefly, swim motility plates (MH broth containing 0.3 % [w/v] agar) were stab inoculated. Swarm motility plates (MH broth containing 0.6 % [w/v] agar) were inoculated by spotting 1 μL of overnight culture. Inoculated plates were then incubated at 21 °C (ambient room temperature) for 8 and 24 hours and 37 °C for 8 hours. The diameter of visible colony spread was measured in mm from three directions and the average value recorded.

### Tolerance to bile salts

The MIC and MBC for sodium deoxycholate (DOC) and sodium cholate was determined for all isolates using the broth dilution method according to Clinical and Laboratory Standard Institute (CLSI) guidelines adapted from the protocol as described previously^53^.

### Bacterial growth curves

The growth of all isolates in LB broth was assessed using a Multiskan FC microplate photometer (Thermo Fisher Scientific) (Supplementary Fig. S3). Measurements were taken every 15 minutes over 24 hours at OD_620_ nm. The instrument was kept at 37 °C with shaking during kinetic intervals.

### Preparation of bacterial inoculum for infection

The MIC and MBC for gentamicin (CN) was determined for all isolates using the broth dilution method according to CLSI guidelines. Susceptibility or resistance was classified according to the lower working concentration of 20 μg/mL used in the gentamicin protection assays as detailed below.

Inoculum stocks for each isolate were prepared by streaking and restreaking individual bacterial isolates onto LB agar. Individual colonies from restreaked isolates were taken and used to inoculate 5 mL LB broth before growth overnight at 37 °C with orbital shaking at 200 RPM. Overnight cultures were centrifuged at 5,500 RCF for 10 minutes. The supernatant was discarded and the bacterial cell pellet resuspended in 5 mL PBS before centrifugation again at 5,500 RCF for 10 minutes. Finally, the pellet was resuspended in 5 mL of PBS (15 % [v/v] glycerol) solution before aliquoting (250 μL/microcentrifuge tube) and freezing at −80 °C. Representative inoculum stocks for each isolate were decimally diluted in PBS and 100 μL aliquots of the dilutions were plated onto LB agar. Agar plates were incubated for 18 hours at 37 °C before enumeration of the CFU.

### *Ex vivo* gentamicin protection assay

The ability to survive and proliferate following phagocytosis by murine RAW 264.7 and human THP-1 macrophages was assessed for all gentamicin susceptible isolates using S. Typhimurium 14028S and ST4/74 as reference strains adapted from protocols as described previously^30,54^.

RAW 264.7 macrophages were grown in antibiotic-free Dulbecco’s Modified Eagle’s Medium (DMEM) (Sigma-Aldrich) supplemented with 10 % [v/v] heat inactivated foetal bovine serum (FBS) and incubated at 37 °C in a humidified atmosphere with 5 % CO_2_. THP-1 monocytes were grown in antibiotic-free RPMI 1640 media (Sigma-Aldrich) supplemented with 10 % [v/v] heat inactivated FBS and incubated at 37 °C in a humidified atmosphere with 5% CO_2_. Cell viability was assessed using trypan blue and a Bio-Rad TC20 automated cell counter.

Cells were subcultured and 1 mL was directly seeded into 24 well plates at a density of 1 × 10^5^ cells/mL per well. THP-1 monocytes were differentiated to adherent macrophages by supplementing media with 20 ng/mL phorbol 12-myristate 13-acetate (PMA) for 5 days.

Prior to infection, inoculum stocks for each bacterial isolate to be assessed were diluted in complete media to 1 × 10^6^ bacteria/mL for a MOI of 10:1 and incubated at 37 °C for 1 hour. Macrophages were washed 3 times with 1 mL Hank’s Balanced Salt Solution (HBSS) before 1 mL of the bacterial suspension prepared as outlined above, was added to each well with 1 mL of complete media being added to uninfected control wells. These 24 well plates were centrifuged at 300 RCF for 5 minutes at room temperature (21 °C) before incubation at 37 °C with 5 % CO_2_ for 1 hour to allow for phagocytosis.

Following phagocytosis, the cells were washed 3 times with 1 mL HBSS. A volume of 1 mL complete media supplemented with 100 μg/mL gentamicin was added to each well before incubation at 37 °C with 5% CO_2_ for 1 hour to kill external bacteria. After 1 hour, cells were washed 3 times with 1 mL HBSS. Another volume of 1 mL of complete media supplemented with 20 μg/mL gentamicin was then added to each well before incubation at 37 °C with 5 % CO_2_ for the desired time points.

Time points were processed by washing the cells 3 times with 1 mL HBSS before 1 mL 1 % [v/v] Triton X-100 PBS solution was added to the infected cells prior to incubation at room temperature for 10 minutes. Lysed supernatants were decimally diluted in PBS and 100 μL aliquots of the dilutions were plated onto LB agar. Agar plates were incubated for 18 hours at 37 °C before enumeration of the CFU.

### Relative viability assay for RAW 264.7 and THP-1 by glucose-6-phosphate dehydrogenase and lactate dehydrogenase

Relative viability of infected macrophages was determined by comparison to uninfected control cells using the Vybrant Cytotoxicity Assay kit (Life Technologies) and Pierce LDH Cytotoxicity Assay kit (Life Technologies) to measure extracellular glucose-6-phosphate dehydrogenase (G6PD) [EC 1.1.1.49] and lactate dehydrogenase (LDH) [EC 1.1.1.27] activity in cell culture supernatants according to manufacturer’s instructions. All samples and standards were assayed in duplicate.

### Cytokine quantification

A panel of proinflammatory cytokines and infection relevant chemokines were quantified from the supernatants of infected RAW 264.7 and THP-1 macrophages (Supplementary Table 10). For RAW 264.7 supernatants, targets were quantified at 0 hours (before infection) and 1, 2, 4, 8 and 24 HPI. For THP-1 supernatants, targets were quantified at 0 hours (before infection) and 1, 2, 4, 8, 24 and 168 HPI.

RAW 264.7 cytokine/chemokine release was measured using a multiplex magnetic bead based kit (Life Technologies) and the Luminex 200 ×MAP platform (Luminex). Similarly, THP-1 cytokine/chemokine release was measured using an electrochemiluminescence based V-PLEX kit (Meso Scale Discovery) and the Sector Imager 2400 platform (Meso Scale Discovery). The measured levels of many targets were below the range of detection in RAW 264.7 supernatant samples and for the purposes of this study were recorded at the lower level of detection for the assay used when comparing to S. Typhimurium ST4/74 infection or infected THP-1 macrophages. Assays were performed according to the manufacturer’s protocol for cell culture supernatant samples. All samples and standards were assayed in duplicate.

### Bacterial whole genome sequencing

Whole genome sequencing was carried out as described previously^55^. Briefly, genomic DNA (gDNA) was purified from overnight cultures of bacterial isolates grown in Trypticase Soy Broth (TSB) (Becton Dickinson) incubated at 37 °C using the DNeasy blood and tissue kit (Qiagen). Libraries were prepared using 1 ng gDNA with the Nextera XT kit (Illumina) and sequenced using the MiSeq platform (Illumina) with a V2 kit (2 × 250 bp).

Three isolates, namely S. Cubana CFSAN002050, S. Tennessee CFSAN001387 and S. Weltevreden CFSAN001415 were further sequenced using the Pacific Biosciences (PacBio) RS II platform. *Salmonella* Cubana CFSAN002050 was sequenced as described previously^56,57^. For S. Tennessee CFSAN001387 and S. Weltevreden CFSAN001415, libraries using 6 μg gDNA were sheared to a size of 10 kb using g-TUBEs (Covaris Inc., Woburn, MA) according to the manufacturer’s instructions. The SMRTbell 10-kb template libraries were constructed using DNA Template Prep Kit 1.0 with the 10-kb insert library protocol (Pacific Biosciences, Menlo Park, CA, USA) and sequenced using the P4-C2 chemistry on 3 single molecule real-time (SMRT) cells with a 240-minute collection protocol along with Stage Start.

### Whole genome assembly and annotation

For Illumina MiSeq data, Jellyfish (version 2.2.6) was used to generate a *k*-mer spectrum before inspecting the quality of the reads using FastQC (version 0.11.5)^58,59^. Error correction was performed using BFC (version r181)^60^. A relaxed sliding window trim for an average Phred quality score of 10 was performed using Trimmomatic (version 0.36) before the genomes were *de novo* assembled with SPAdes (version 3.7.1) using the default *k*-mer size selection for 250 bp reads and the automatic coverage cutoff threshold^61,62^. The quality of the subsequent assemblies was assessed using Bandage (version 0.8.0) and QUAST (version 4.1)^63,64^. Contigs were excluded from the assembly if they were shorter than 200 bp.

Analysis of the PacBio data was implemented using SMRT Analysis 2.3.0. The best *de novo* assembly was established with PacBio Hierarchical Genome Assembly Process (HGAP 3.0) program using the continuous-long-reads from the three SMRT cells. The assembly outputs from HGAP produced circular contiguous sequences with overlapping regions at the end that can be identified using dot plots in Gepard (version 1.40)^65^. Genomes were checked manually for even sequencing coverage. Afterwards the interim consensus sequence was used to determine the final consensus and accuracy scores using Quiver consensus algorithm^66^.

All sequences and assemblies are publicly available with accession numbers provided (Table 1) and are submitted for annotation using the NCBI Prokaryotic Genome Automatic Annotation Pipeline (PGAAP)^67^.

### Sequence analysis

*<u>S</u>almonella* <u>P</u>athogenicity Island (SPI) gene content was compared to S. Typhimurium ST4/74 for all isolates (Fig. 5 and Supplementary Fig. S8). Homologous amino acid sequences for each protein were identified and generated using BLAST+ (version 2.3.0) and Biopython (version 1.66)^68–70^. Amino acid sequence similarity was assessed using a Needleman-Wunsch global alignment through the EMBOSS analysis software (version 6.6.0)^71^.

### Pan-genome analysis

To limit bias among different annotation tools for downstream analyses, additional annotation of all isolates included in this study was performed using Prokka (version 1.1 1)^72–79^. Presence or absence of protein sequences from all strains was determined using the pan-genome pipeline Roary (version 3.6.1) (Supplementary Fig. S9b and Supplementary Fig. S9c)^80–83^. Visualisation of the pan-genome data was performed using Anvi’o (version 1.2.3) (Supplementary Fig. S9a)^84,85^.

### Statistics

The R statistical computing environment was used for all statistical analyses^86,87^. Multiple comparisons for normally distributed data were performed by one-way ANOVA where appropriate and *post hoc* analysis of significance was inspected by Tukey’s range test.

### Code availability

Data and code to reproduce the analyses and manuscript figures are available^88^.

## Acknowledgements

Daniel Hurley is supported by the Wellcome Trust Computational Infection Biology PhD Programme (Grant reference: 099837/Z/12/Z).

## Author information

## Contributions

D.H. and M.M. designed experiments. D.H. and M.H. performed experiments and analysed data. T.M. performed experiments. D.H. and S.F. wrote the manuscript. All authors discussed the results and contributed to the revision of the manuscript.

## References

1. Kirk M. D. et al. World Health Organization Estimates of the Global and Regional Disease Burden of 22 Foodborne Bacterial, Protozoal, and Viral Diseases, 2010: A Data Synthesis. PLoS Med 12, e1001921 (2015).

2. Crump J. A., Luby S. P. & Mintz E. D. The global burden of typhoid fever. Bull. World Health Organ. 82, 346–353 (2004).

3. Majowicz S. E. et al. The Global Burden of Nontyphoidal Salmonella Gastroenteritis. Clin. Infect. Dis. 50, 882–889 (2010).

4. Baumler A. J., Tsolis R. M., Ficht T. A. & Adams L. G. Evolution of Host Adaptation inSalmonella enterica. Infect. Immun. 66, 4579–4587 (1998).

5. Barton Behravesh C. et al. 2008 outbreak of Salmonella Saintpaul infections associated with raw produce. N. Engl. J. Med. 364, 918–927 (2011).

6. Riesenberg-Wilmes, M. R., Bearson B., Foster J. W. & Curtis R. Role of the acid tolerance response in virulence of Salmonella typhimurium. Infect. Immun. 64, 1085–1092 (1996).

7. Prouty A. M., Brodsky I. E., Falkow S. & Gunn J. S. Bile-salt-mediated induction of antimicrobial and bile resistance in Salmonella typhimurium. Microbiol. Read. Engl. 150, 775–783 (2004).

8. Haraga A., Ohlson M. B. & Miller S. I. Salmonellae interplay with host cells. Nat. Rev. Microbiol. 6, 53–66 (2008).

9. Juhas M. et al. Genomic islands: tools of bacterial horizontal gene transfer and evolution. Fems Microbiol. Rev. 33, 376–393 (2009).

10. Steele-Mortimer, O. The Salmonella-containing Vacuole – Moving with the Times. Curr. Opin. Microbiol. 11, 38–45 (2008).

11. Coburn B., Grassl G. A. & Finlay B. B. Salmonella, the host and disease: a brief review. Immunol. Cell Biol. 85, 112–118 (2006).

12. Hurley D., McCusker M. P., Fanning, S. & Martins, M. Salmonella-host interactions — modulation of the host innate immune system. Microb. Immunol. 5, 481 (2014).

13. Zhang S. et al. Molecular Pathogenesis of Salmonella enterica Serotype Typhimurium-Induced Diarrhea. Infect. Immun. 71, 1–12 (2003).

14. Kroger C. et al. The transcriptional landscape and small RNAs of Salmonella enterica serovar Typhimurium. Proc. Natl. Acad. Sci. 109, E1277–E1286 (2012).

15. Kroger C. et al. An Infection-Relevant Transcriptomic Compendium for Salmonella enterica Serovar Typhimurium. Cell Host Microbe 14, 683–695 (2013).

16. Sabbagh S. C., Lepage C., McClelland M. & Daigle F. Selection of Salmonella enterica Serovar Typhi Genes Involved during Interaction with Human Macrophages by Screening of a Transposon Mutant Library. PLoS ONE 7, e36643 (2012).

17. Srikumar S. et al. RNA-seq Brings New Insights to the Intra-Macrophage Transcriptome of Salmonella Typhimurium. PLoS Pathog 11, e1005262 (2015).

18. Westermann A. J. et al. Dual RNA-seq unveils noncoding RNA functions in host-pathogen interactions. Nature 529, 496–501 (2016).

19. Olsen S. J. et al. The Changing Epidemiology of Salmonella: Trends in Serotypes Isolated from Humans in the United States, 1987-1997. J. Infect. Dis. 183, 753–761(2001).

20. Guibourdenche M. et al. Supplement 2003-2007 (No. 47) to the White-Kauffmann-Le Minor scheme. Res. Microbiol. 161, 26–29 (2010).

21. Achtman M. et al. Multilocus Sequence Typing as a Replacement for Serotyping in Salmonella enterica. PLoS Pathog 8, e1002776 (2012).

22. Makendi C. et al. A Phylogenetic and Phenotypic Analysis of Salmonella enterica Serovar Weltevreden, an Emerging Agent of Diarrheal Disease in Tropical Regions. PLoS Negl Trop Dis 10, 1–19 (2016).

23. Lockman H. A. & Curtiss R. Salmonella typhimurium mutants lacking flagella or motility remain virulent in BALB/c mice. Infect. Immun. 58, 137–143 (1990).

24. Li J. et al. Evolutionary origin and radiation of the avian-adapted non-motile salmonellae. J. Med. Microbiol. 38, 129–139 (1993).

25. Toguchi A., Siano M., Burkart M. & Harshey R. M. Genetics of Swarming Motility in Salmonella enterica Serovar Typhimurium: Critical Role for Lipopolysaccharide. J. Bacteriol. 182, 6308–6321 (2000).

26. Harshey R. M. Bacterial Motility on a Surface: Many Ways to a Common Goal. Annu. Rev. Microbiol. 57, 249–273 (2003).

27. Lathrop S. K. et al. Replication of Salmonella Typhimurium in Human Monocyte-Derived Macrophages. Infect. Immun. IAI.00033-15 (2015). doi: 10.1128/IAI.00033-15

28. Schwan W. R., Huang X.-Z., Hu L. & Kopecko D. J. Differential Bacterial Survival, Replication, and Apoptosis-Inducing Ability of Salmonella Serovars within Human and Murine Macrophages. Infect. Immun. 68, 1005–1013 (2000).

29. Vaudaux P. & Waldvogel F. A. Gentamicin antibacterial activity in the presence of human polymorphonuclear leukocytes. Antimicrob. Agents Chemother. 16, 743–749 (1979).

30. Steele-Mortimer, O., Meresse S., Gorvel, J.-P., Toh, B.-H. & Finlay B. B. Biogenesis of Salmonella typhimurium-containing vacuoles in epithelial cells involves interactions with the early endocytic pathway. Cell. Microbiol. 1, 33–49 (1999).

31. Steele-Mortimer, O. et al. The invasion-associated type III secretion system of Salmonella enterica serovar Typhimurium is necessary for intracellular proliferation and vacuole biogenesis in epithelial cells. Cell. Microbiol. 4, 43–54 (2002).

32. Malik-Kale P., Winfree S. & Steele-Mortimer, O. The Bimodal Lifestyle of Intracellular Salmonella in Epithelial Cells: Replication in the Cytosol Obscures Defects in Vacuolar Replication. PLoS ONE 7, e38732 (2012).

33. Braukmann M., Methner U. & Berndt A. Immune Reaction and Survivability of Salmonella Typhimurium and Salmonella Infantis after Infection of Primary Avian Macrophages. PLoS ONE 10, e0122540 (2015).

34. Modi W. S. & Yoshimura T. Isolation of novel GRO genes and a phylogenetic analysis of the CXC chemokine subfamily in mammals. Mol. Biol. Evol. 16, 180–193 (1999).

35. McWhorter A. R. & Chousalkar K. K. Comparative phenotypic and genotypic virulence of Salmonella strains isolated from Australian layer farms. Food Microbiol. 6, 12 (2015).

36. Dhanani A. S. et al. Genomic Comparison of Non-Typhoidal Salmonella enterica Serovars Typhimurium, Enteritidis, Heidelberg, Hadar and Kentucky Isolates from Broiler Chickens. PLoS ONE 10, e0128773 (2015).

37. Ehrbar K., Friebel A., Miller S. I. & Hardt, W.-D. Role of the Salmonella Pathogenicity Island 1 (SPI-1) Protein InvB in Type III Secretion of SopE and SopE2, Two Salmonella Effector Proteins Encoded Outside of SPI-1. J. Bacteriol. 185, 6950–6967 (2003).

38. Hapfelmeier S. et al. The Salmonella pathogenicity island (SPI)-2 and SPI-1 type III secretion systems allow Salmonella serovar typhimurium to trigger colitis via MyD88-dependent and MyD88-independent mechanisms. J. Immunol. Baltim. Md 1950 174, 1675–1685 (2005).

39. Collier-Hyams L. S. et al. Cutting Edge: Salmonella AvrA Effector Inhibits the Key Proinflammatory Anti. Apoptotic NF-kB Pathway. J. Immunol. 169, 2846–2850 (2002).

40. Liao A. P. et al. Salmonella Type III Effector AvrA Stabilizes Cell Tight Junctions to Inhibit Inflammation in Intestinal Epithelial Cells. PLoS ONE 3, e2369 (2008).

41. Lin S. L., Le T. X. & Cowen D. S. SptP, a Salmonella typhimurium type III-secreted protein, inhibits the mitogen-activated protein kinase pathway by inhibiting Raf activation. Cell. Microbiol. 5, 267–275 (2003).

42. Waterman S. R. & Holden, D. W. Functions and effectors of the Salmonella pathogenicity island 2 type III secretion system. Cell. Microbiol. 5, 501–511 (2003).

43. Figueira R. & Holden D. W. Functions of the Salmonella pathogenicity island 2 (SPI-2) type III secretion system effectors. Microbiol. Read. Engl. 158, 1147–1161 (2012).

44. Freeman J. A., Rappl C., Kuhle V., Hensel M. & Miller S. I. SpiC Is Required for Translocation of Salmonella Pathogenicity Island 2 Effectors and Secretion of Translocon Proteins SseB and SseC. J. Bacteriol. 184, 4971–4980 (2002).

45. Nikolaus T. et al. SseBCD Proteins Are Secreted by the Type III Secretion System of Salmonella Pathogenicity Island 2 and Function as a Translocon. J. Bacteriol. 183, 6036–6045 (2001).

46. Gerlach R. G. et al. Salmonella Pathogenicity Island 4 encodes a giant non-fimbrial adhesin and the cognate type 1 secretion system. Cell. Microbiol. 9, 1834–1850 (2007).

47. Gerlach R. G., Jackel D., Geymeier N. & Hensel M. Salmonella Pathogenicity Island 4-Mediated Adhesion Is Coregulated with Invasion Genes in Salmonella enterica. Infect. Immun. 75, 4697–4709 (2007).

48. Morgan E., Bowen A. J., Carnell S. C., Wallis T. S. & Stevens M. P. SiiE Is Secreted by the Salmonella enterica Serovar Typhimurium Pathogenicity Island 4-Encoded Secretion System and Contributes to Intestinal Colonization in Cattle. Infect. Immun. 75, 1524–1533 (2007).

49. Wood M. W. et al. Identification of a pathogenicity island required for Salmonella enteropathogenicity. Mol. Microbiol. 29, 883–891 (1998).

50. National Salmonella Surveillance | National Surveillance | CDC. Available at: http://www.cdc.gov/nationalsurveillance/salmonella-surveillance.html.

51. Allard M. W. et al. the PRACTICAL value of Food Pathogen Traceability through BUILDING a Whole-Genome Sequencing Network and database. J. Clin. Microbiol. (2016). doi:10.1128/JCM.00081-16

52. Martins M. et al. Evidence of Metabolic Switching and Implications for Food Safety from the Phenome(s) of Salmonella enterica Serovar Typhimurium DT104 Cultured at Selected Points across the Pork Production Food Chain. Appl. Environ. Microbiol. 79, 5437–5449 (2013).

53. Hernandez S. B., Cota I., Ducret A., Aussel L. & Casadesus J. Adaptation and Preadaptation of Salmonella enterica to Bile. PLoS Genet 8, e1002459 (2012).

54. Drecktrah D., Knodler L. A., Ireland R. & Steele-Mortimer, O. The mechanism of Salmonella entry determines the vacuolar environment and intracellular gene expression. Traffic Cph. Den. 7, 39–51 (2006).

55. Toro M. et al. Draft Genome Sequences of 33 Salmonella enterica Clinical and Wildlife Isolates from Chile. Genome Announc. 3, e00054–15 (2015).

56. Hoffmann M. et al. First Fully Closed Genome Sequence of Salmonella enterica subsp. enterica Serovar Cubana Associated with a Food-Borne Outbreak. Genome Announc. 2, e01112–14 (2014).

57. Pirone-Davies, C. et al. Genome-Wide Methylation Patterns in Salmonella enterica Subsp. enterica Serovars. PLoS ONE 10, e0123639 (2015).

58. Simon, A. FastQC: A quality control tool for high throughput sequence data. (2010).

59. Margais, G. & Kingsford C. A fast, lock-free approach for efficient parallel counting of occurrences of k-mers. Bioinformatics 27, 764–770 (2011).

60. Li, H. BFC: correcting Illumina sequencing errors. Bioinformatics 31, 2885–2887 (2015).

61. Bankevich, A. et al. SPAdes: A New Genome Assembly Algorithm and Its Applications to Single-Cell Sequencing. J. Comput. Biol. 19, 455–477 (2012).

62. Bolger A. M., Lohse, M. & Usadel, B. Trimmomatic: a flexible trimmer for Illumina sequence data. Bioinformatics 30, 2114–2120 (2014).

63. Gurevich, A., Saveliev, V., Vyahhi, N. & Tesler, G. QUAST: quality assessment tool for genome assemblies. Bioinformatics 29, 1072–1075 (2013).

64. Wick R. R., Schultz M. B., Zobel, J. & Holt K. E. Bandage: interactive visualization of de novo genome assemblies. Bioinformatics 31, 3350–3352 (2015).

65. Krumsiek J., Arnold R. & Rattei T. Gepard: a rapid and sensitive tool for creating dotplots on genome scale. Bioinformatics 23, 1026–1028 (2007).

66. Chin, C.-S. et al. Nonhybrid, finished microbial genome assemblies from long-read SMRT sequencing data. Nat. Methods 10, 563–569 (2013).

67. Klimke W. et al. The National Center for Biotechnology Information’s Protein Clusters Database. Nucleic Acids Res. 37, D216–D223 (2009).

68. Altschul S. F., Gish W., Miller W., Myers E. W. & Lipman D. J. Basic local alignment search tool. J. Mol. Biol. 215, 403–410 (1990).

69. Camacho C. et al. BLAST+: architecture and applications. BMC Bioinformatics 10, 421 (2009).

70. Cock P. J. A. et al. Biopython: freely available Python tools for computational molecular biology and bioinformatics. Bioinformatics 25, 1422–1423 (2009).

71. Rice P., Longden I. & Bleasby A. EMBOSS: The European Molecular Biology Open Software Suite. Trends Genet. 16, 276–277 (2000).

72. Laslett D. & Canback B. ARAGORN, a program to detect tRNA genes and tmRNA genes in nucleotide sequences. Nucleic Acids Res. 32, 11–16 (2004).

73. Hyatt D. et al. Prodigal: prokaryotic gene recognition and translation initiation site identification. BMC Bioinformatics 11, 119 (2010).

74. Finn R. D., Clements J. & Eddy S. R. HMMER web server: interactive sequence similarity searching. Nucleic Acids Res. 39, W29–W37 (2011).

75. Petersen T. N., Brunak S., von Heijne G. & Nielsen H. SignalP 4.0: discriminating signal peptides from transmembrane regions. Nat. Methods 8, 785–786 (2011).

76. Nawrocki E. P. & Eddy S. R. Infernal 1.1: 100-fold faster RNA homology searches. Bioinformatics 29, 2933–2935 (2013).

77. Seemann T. Prokka:rapid prokaryotic genome annotation. Bioinformatics 30, 2068–2069 (2014).

78. tseemann/barrnap. GitHub Available at: https://github.com/tseemann/barrnap.

79. ctSkennerton/minced. GitHub Available at: https://github.com/ctSkennerton/minced.

80. Enright A. J., Dongen S. V. & Ouzounis C. A. An efficient algorithm for large-scale detection of protein families. Nucleic Acids Res. 30, 1575–1584 (2002).

81. Li W. & Godzik A. Cd-mhit: a fast program for clustering and comparing large sets of protein or nucleotide sequences. Bioinformatics 22, 1658–1659 (2006).

82. Quinlan A. R. & Hall I. M. BEDTools: a flexible suite of utilities for comparing genomic features. Bioinformatics 26, 841–842 (2010).

83. Page A. J. et al. Roary: rapid large-scale prokaryote pan genome analysis. Bioinformatics 31, 3691–3693 (2015).

84. Price M. N., Dehal P. S. & Arkin A. P. FastTree 2 – Approximately Maximum-Likelihood Trees for Large Alignments. PLOS ONE 5, e9490 (2010).

85. Eren A. M. et al. Anvi’o: an advanced analysis and visualization platform for ‘omics data. PeerJ 3, e1319 (2015).

86. Wickham H. ggplot2: Elegant Graphics for Data Analysis. (Springer-Verlag New York, 2009).

87. R Core Team. R: A Language and Environment for Statistical Computing. (R Foundation for Statistical Computing, 2016).

88. danieljhurley/atypical-salmonella. GitHub Available at: https://github.com/danieljhurley/atypical-salmonella.

